# Growth and development of trabecular structure in the calcaneus of Japanese macaques (*Macaca fuscata*) reflects locomotor behavior, life-history, and neuromuscular development

**DOI:** 10.1101/2021.10.07.463526

**Authors:** Jaap P.P. Saers, Adam D. Gordon, Timothy M. Ryan, Jay T. Stock

## Abstract

We aim to broaden the analysis of bone structure by suggesting a new way to incorporate the interactions between behavior, neuromuscular development, and life-history. We examine the associations between these variables and age-related variation in trabecular structure in the calcaneus of Japanese macaques (*Macaca fuscata*). If skeletal markers linking these variables can be established, our inferences of the biology and behavior of fossil species would be significantly improved. We µCT scanned the calcaneus in a cross-sectional sample of 36 juveniles aged between 0 and 7 years old and 5 adults at the Primate Research Institute, Japan. We calculated whole bone averages of standard trabecular properties and generated whole-bone morphometric maps of bone volume fraction and Young’s modulus. Trabecular structure is increasingly heterogeneous in older individuals. BV/TV decreases during the first month of life and increases afterwards, coinciding with the onset of independent locomotion. At birth, primary Young’s modulus is oriented orthogonal to the ossification center, but after locomotor onset bone structure becomes stiffest in the direction of joint surfaces and muscle attachments. Age-related variation in bone volume fraction is best predicted by an interaction between neuromaturation, body mass, and locomotor independence. Results support the common assumption that trabecular structure dynamically adapts to novel joint loading conditions during ontogeny. The timing of independent locomotion, body size, and neuromuscular development, are all correlated to age-related variation in the trabecular structure of the macaque calcaneus. The causal mechanisms behind the observed patterns cannot be directly inferred from our cross-sectional study. If the model presented in this paper holds up under longitudinal experimental conditions, trabecular structure can be used both to infer behavior from fossil morphology and to serve as a valuable proxy for neuromuscular maturation and life history events like locomotor onset and the achievement of an adult-like gait.

## Introduction

Bone morphology is partially determined by genetic processes regulating growth and development, and partially by bone cells sensing and responding to their mechanical environment (Huiskes et al., 2000; Carter and Beaupré, 2001; Currey, 2002). The structure of spongy (trabecular) bone found inside bones is thought to be particularly responsive to mechanical stimuli (Carter et al., 1996; Carter and Beaupré, 2001; Kivell, 2016). The link between mechanical loading and the three-dimensional structure of trabecular bone allows locomotor and postural behavior to be reconstructed in fossil taxa (Ryan and Ketcham, 2002; Skinner et al., 2015; Kivell, 2016; Stephens et al., 2016; Zeininger et al., 2016; Bishop et al., 2018; Ryan et al., 2018; Bardo et al., 2020; Dunmore et al., 2020). To understand how variation in trabecular structure arises within and between species, it is imperative to understand how it forms during growth and development (Ryan and Krovitz, 2006; Gosman and Ketcham, 2009; Ryan et al., 2017; Saers et al., 2020). Indeed, alterations to ontogenetic trajectories are the principal ways in which evolutionary changes in life history and morphology occur (Gould, 1977; Hallgŕmsson and Hall, 2005; Kardong, 2018; Woronowicz and Schneider, 2019). Recent methodological and technological advances in the analysis of trabecular bone structure have opened new possibilities for studying the development of trabecular bone structure (Gross et al., 2014; DeMars et al., 2020). Here we apply these new techniques to analyse the ontogeny of trabecular structure in the calcaneus of Japanese macaques (*Macaca fuscata*).

The musculoskeletal system undergoes striking changes throughout growth and development. Movements starting in utero and continuously changing throughout development, generate loads that shape bone morphology into the general adult form (Carter and Beaupré, 2001). These processes are essential for generating the adult morphology that is required for typical species-specific gait and posture (Tardieu, 1999). Mammalian locomotion usually develops in a stereotypical sequence of events that dramatically change how their skeletons are loaded (Doran, 1997; Lacquaniti et al., 2012; Sarringhaus et al., 2014). Changes in loading direction, magnitude, and frequency (Rubin et al., 2002; Sugiyama et al., 2010; Barak et al., 2011) alter the trabecular structure during development, so that the structure provides a ‘functional record’ of behavioural changes throughout development (Wolschrijn and Weijs, 2004; Pontzer et al., 2006; Ryan and Krovitz, 2006; Gosman and Ketcham, 2009; Barak et al., 2011; Ryan et al., 2017; Tsegai et al., 2018; Saers et al., 2020, 2021). Although the study of trabecular bone development is in its infancy, previous research on the humans (Ryan and Krovitz, 2006; Cunningham and Black, 2009; Gosman and Ketcham, 2009; Reissis and Abel, 2012; Acquaah et al., 2015; Raichlen et al., 2015; Ryan et al., 2017; Colombo et al., 2019; Saers et al., 2020), great apes (Tsegai et al., 2018; Ragni, 2020), and other mammals (Tanck et al., 2001; Wolschrijn and Weijs, 2004; Gorissen et al., 2016) suggests a strong link between changes in loading conditions as gait develops and responses in trabecular structure.

### Trabecular bone ontogeny

The connections between trabecular morphology and habitual loading patterns has been demonstrated experimentally (Barak et al., 2011), but trabecular bone ontogeny is still not yet thoroughly understood, particularly how the degree of plasticity versus genetic canalization varies throughout development and into adulthood (Cunningham and Black, 2009; Reissis and Abel, 2012; Raichlen et al., 2015; Gorissen et al., 2016; Ryan et al., 2017; Saers et al., 2020). Bone growth occurs via the transformation of growth plate cartilage into bone through a series of cell and matrix changes (Byers et al., 2000; Parfitt et al., 2000; Burr and Organ, 2017). The transformation from growth plate cartilage to trabecular bone is similar among mammals, indicating a highly conserved process (Frost and Jee, 1994; Byers et al., 2000). This process sets up a basic trabecular structure which can later be modified through metabolic and mechanical factors. Trabecular bone is laid out orthogonal to the growth plate in a dense and anisotropic structure which is later refined into bone- and species-specific heterogeneous adult states. Frost and Jee (1994) argue that the effects of mechanical usage during periods of rapid bone growth in early ontogeny explain many of the features observed during the ossification process. They propose that mechanical strain is the controlling mechanism for endochondral ossification, in which the underloaded elements of the dense bone structure during the first years of life are removed and bone is added in strained areas, resulting in a mechanically adapted state (Frost & Jee, 1994). This model correctly predicts observations of bone loss at early stages of ontogeny and explains it as the result of the removal of redundant material below a certain strain threshold (Carter et al., 1991, 1996; Carter and Beaupré, 2001; Frost, 2003; Pivonka et al., 2018).

### The brain-bone connection

Brains and trabecular bone have more in common than one might initially think. Both are made up of complex, interconnected three dimensional structures and broadly share developmental patterns. At birth, both trabeculae and neurons are overproduced. Structures are refined to a more heterogeneous state during ontogeny through modeling in bone and synaptic pruning in neurons. This happens under the influence of some input, presumably mechanical in terms of trabecular bone (Huiskes et al., 2000; Carter and Beaupré, 2001), and through neural activity in the brain (Shatz, 1990; Sakai, 2020). In both cases, there is a long history of debate as to how much of its respective morphology is genetically canalized versus plastic in response to its environment, i.e. nature versus nurture, and in both cases the consensus is “both”. While starting with an excess of connections to remove many of them later may seem inefficient, the result is a state that is adapted to an individual’s specific environment. Indeed, this process is so efficient that it is found in many other tissues as well including connective tissues like ligaments and tendons (Grinnell, 2000) to the nervous system (Sakai, 2020).

The patterns of the growth and development of trabecular structure reviewed above are consistent with a model where a generalized trabecular structure is formed by dynamic adaptation to local, bone- and region-specific loading patterns. These loading patterns are generated by neural circuits that develop in parallel to increases in physical size and weight of a growing organism (Forssberg, 1985). Locomotor patterns are transformed from an immature state to increasingly adult-like patterns during development. During the early ontogeny of gait, infants are mainly focused on minimizing the risk of falling. When individuals increase in strength, stability improves and postural constraints are reduced (Vaughan et al., 2003). It is thought that development subsequently proceeds to select the most optimal neural networks (Forssberg, 1999), resulting in a reduction in the variability of muscle activation and co-contraction, and the adult gait pattern emerges (Okamoto et al., 2003). If trabecular structure is a reliable reflection of gait mechanics, then changes in trabecular structure during growth should reflect gait mechanics, which in turn reflects the degree of neurological maturation of locomotion, as well as an animal’s degree of precociality. If this link can be demonstrated, then trabecular structure could be a valuable proxy for neuromuscular maturation in fossil species.

Across mammals (Garwicz et al., 2009) and birds (Iwaniuk and Nelson, 2003), adult brain size strongly predicts time to locomotor onset after conception. In addition, the onset of walking is strongly correlated with the timing of several important aspects of brain development. In humans, locomotion is not just a developmental precursor to numerous psychological changes but plays a causal role in their formation (Campos et al., 2000; Uchiyama et al., 2008; Anderson et al., 2013; Dahl et al., 2013). The onset of human independent locomotion is followed by a revolution in perception-action coupling, spatial cognition, memory, and social and emotional development (Anderson et al., 2013). Research indicates that neural function and structure reciprocally influence one another throughout development (Campos et al., 2000; Anderson et al., 2013), placing the activity of locomotor development in the center of development, rather than being just a consequence of neural maturation. In other words, the onset of independent locomotion is an important life-history event related to adult brain size and the timing of neuromuscular development. If we can detect bony markers of locomotor development, this would be able to provide a unique insight into fossil locomotion as well as aspects of life history (Zihlman, 1992).

### Locomotor development in Japanese macaques

The basic locomotor characteristics of Japanese macaques appear in the first two months after birth (Torigoe, 1984; Nakano, 1996; Dunbar and Badam, 1998; Kimura, 2000). Newborns cannot walk, stand, or sit on their own and reflexively hang on to their mother for transport. Initial, somewhat poorly coordinated quadrupedal movements emerge near the end of the first month (Torigoe, 1984; Nakano, 1996). Macaques begin to locomote primarily by walking in both diagonal and lateral sequences, followed after four weeks by occasionally running and trotting (Nakano, 1996).

Independent locomotion away from the mother becomes regular after two months of age. Coordinated walking appears after three months. Prior to this, their style of walking is limited by the immature development of their musculoskeletal system (Nakano, 1996). Locomotion becomes increasingly refined and independent throughout the first year of life (Dunbar and Badam, 1998). Locomotor activities include unskilled locomotion between one and six months when monkeys still frequently lose their footing. After six months the macaques are skilled at both terrestrial and arboreal locomotion (Torigoe, 1984; Nakano, 1996; Kimura, 2000).

Between the age of 1 to 3-4 years the macaques enter the juvenile phase which contains the most diverse range of posture and locomotion. The juveniles have a well-developed musculoskeletal system which, combined with a small body size, enables juvenile macaques to exploit terrestrial and arboreal environments to their fullest potential (Dunbar and Badam, 1998). After the juvenile phase Macaques are considered adults but they continue to grow in size, albeit at a decreasing rate, until around ten years of age (Hamada et al., 2004). The postural and locomotor repertoires of adults are reduced compared to juveniles due to increases in body size. The largest reduction is among play behaviors in the small-branch setting of trees, and below-branch postures and locomotion disappear (Dunbar and Badam, 1998).

### Aims

If trabecular bone markers of behavior, neuromaturation, and life history variables such as the onset of independent locomotion can be established, reconstructions of fossil behavior would be substantially improved. Our aim is to first document how trabecular structure of the calcaneus of Japanese macaques varies with age and body mass. We then test whether landmark events in the development of locomotion (independent locomotion, achievement of adult-like locomotor repertoires) in macaques coincide with clear signals in the trabecular structure. We do this by analyzing whole-bone averages of standard trabecular properties (Table 1) as well as regional variation in the distribution of these properties throughout the calcaneus. Additionally, we aim to broaden the analysis of bone structure beyond pure mechanics by proposing a new way to incorporate the interactions between behavior and neuromuscular development, body size, and life-history.

**Table 1.** Definitions of trabecular bone properties using in this study.

| Measurement | Abv. | Units | Description |
| --- | --- | --- | --- |
| Bone volume fraction | BV/TV | - | Ratio of bone volume to total volume of interest |
| Trabecular thickness | Tb.Th | mm | Average trabecular strut thickness |
| Trabecular spacing | Tb.Sp | mm | Average distance between struts |
| Degree of anisotropy | DA | - | Extent to which trabeculae are oriented in a particular direction. Ranging from isotropic (0) to anisotropic (1). |
| Primary Young's modulus | E | MPa | The tensile stiffness of a solid material in the direction in which it is stiffest measured in a sphere |

### Hypotheses

If the development of trabecular structure is largely or partially mediated through mechanics (Huiskes et al., 2000; Carter and Beaupré, 2001) rather than genetic programming (Lovejoy et al., 2003), one would predict that a lack of locomotor-related loading during the first month would lead to bone resorption while increases in loading after the onset of locomotion should result in bone formation (Frost, 2003; Pivonka et al., 2018). Japanese macaques start to locomote somewhat regularly in the second month of life and locomotor independence from the mother is established after two months of age (Torigoe, 1984; Nakano, 1996; Kimura, 2000). New mechanical stimulation after the onset of locomotion combined with increases in body size, are predicted to initiate reorganization of the trabecular architecture throughout the calcaneus. Redundant trabeculae are expected to be removed, while trabeculae oriented in directions involved in the distribution of loads associated with locomotion are preserved or enlarged, resulting in a reorganization of the primary direction of bone stiffness, and increases in bone volume fraction and average trabecular thickness. After the onset of locomotion, trabeculae are expected to increase in regional variation in the amount of bone, bone stiffness, and average orientation of trabeculae. The highest BV/TV is expected to be found where loads are applied to the calcaneus, including under joint surfaces (posterior talar facet and calcaneocuboid joint) and the attachment sites of the Achilles tendon and the plantar ligaments (Giddings et al., 2000; Saers et al., 2020).

We predict the following events to invoke the following associated morphological signals:

1. **Onset of locomotion**: the slope of BV/TV with age should shift from negative to positive, trabeculae thicken, primary stiffness changes direction (Barak et al., 2011; Ryan et al., 2017; Saers et al., 2020).
2. **Appearance of adult-like locomotor repertoire:** the increase in BV/TV should level off and follow a more shallow slope due to allometric scaling with still increasing body size (Doube et al., 2011; Ryan and Shaw, 2013; Saers et al., 2019; Mulder et al., 2020).
3. **Neuromuscular maturation of gait:** age-related variation in loading patterns is generated by neural circuits that develop in parallel to increases in physical size and weight of a growing organism. As such, trabecular properties should be predicted by an interaction between body mass and neuromaturation. Here we use the percentage of adult brain size for age as a proxy for neuromuscular maturation.

## Materials and methods

### Sample

We µCT scanned the calcaneus from the skeletal remains of 36 juvenile male Japanese macaques (*Macaca fuscata fuscata*) from a colony housed in a large open air enclosure at the Primate Research Institute, University of Kyoto (PRI) (Torigoe, 1984). Individuals are of known age and body weight at death and in most cases parental lineage is recorded. We also include five wild adult males from the PRI skeletal collection of unknown precise age but whose body weight at death was recorded. Specimens were CT scanned at the PRI with a SkyScan1275 µCT scanner at 95kV and 95µA for 2400 projections with an exposure of 0.216 seconds. CT scans were saved as 16bit tiff stacks with isotropic voxel dimensions between 16 and 22µm depending on bone size. We tested for potential effects of variation in voxel dimensions by artificially reducing the voxel size from 16 to 22µm in Amira 6.7.0 (Thermo Fisher Scientific) using the resample module with a Lanczos filter. Least-squares linear regression analysis was run for each variable to test for significant influences of voxel size on trabecular properties. Downsampling individuals from 16 to 22 µm yielded identical results, indicating the voxel size differences do not affect the results. We calculated percentage of adult body mass for age in *Macaca fuscata* based on data from Hamada (1994) and percentage brain volume for age in by combining data from two studies (DeSilva and Lesnik, 2008; Van Minh and Hamada, 2017) and dividing by the average adult brain volume (105.6 cm^3^, DeSilva and Lesnik, 2008). When then fit the following curve to predict percentage of adult brain volume for age in months: 0.7417*age^0.0681^.

### Calculation of whole bone average trabecular properties

The three-dimensional structure of trabecular bone was quantified using standard trabecular properties (Table 1). Tiff stacks were segmented into bone and non-bone using the MIA clustering algorithm (Dunmore et al., 2018) and then imported into ORS Dragonfly for whole bone quantification of trabecular properties (Object Research Systems (ORS) Inc, Montreal, Canada, www.theobjects.com/dragonfly). We calculated the average bone volume fraction (BV/TV), trabecular thickness (Tb.Th), Trabecular spacing (Tb.Sp), and degree of anisotropy (DA). Degree of anisotropy was calculated using the mean intercept length method.

### 3D mapping of trabecular structure

Segmented scans were categorized into three regions (cortex, trabeculae, and internal region of the bone) using Medtool 4.0 (www.dr-pahr.at, Figure 1). Morphometric maps of BV/TV and primary Young’s modulus were generated following Gross et al. (2014). A three-dimensional (3D) tetrahedral mesh was created of the internal region of the bone using CGAL (http://www.cgal.org). A mesh size of 0.6 mm was used. A 7 mm background grid was applied in three dimensions to the trabecular, and BV/TV and Young’s modulus were quantified at each node of the background grid using a 3.5 mm sampling sphere. The values from each sampling sphere were interpolated and applied to elements of the 3D tetrahedral mesh to generate morphometric maps.

**Figure 1.**
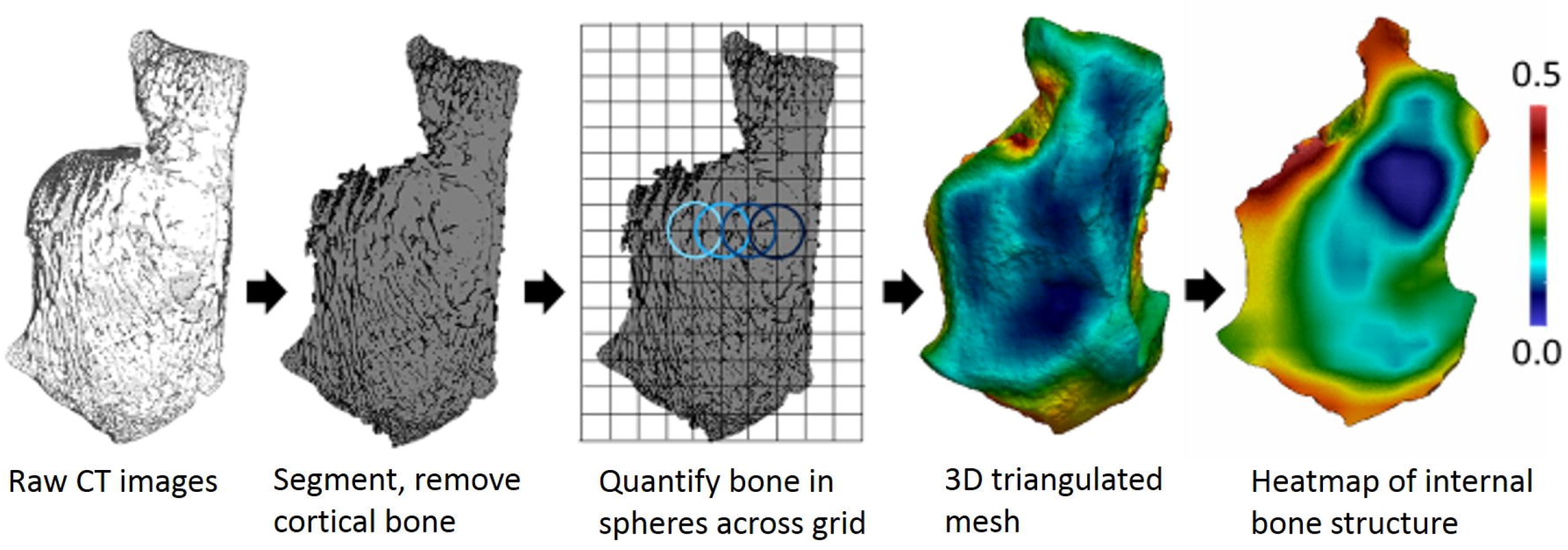
Medtool workflow example for a calcaneus.

### Statistical analysis

Linear regressions and interactions between trabecular properties, age, and body weight were performed in R version 4.0.2 (R Core Team, 2019). Alpha level was set to 0.05 for all statistical tests. When comparing various regression models, the model with the lowest Akaike Information Criterion (AIC; Akaike, 1974) and highest R^2^ was chosen as indicating the highest model quality. Adding additional variables to a regression always increases the fit (R^2^) due to spurious correlations, this process is called overfitting and causes the model to learn too much from the data, resulting in poor predictive power for non-measured samples. AIC measures the degree to which a model is overfit with lower values indicating a greater model quality (Akaike, 1974; McElreath, 2015). In addition to regular linear regression we run piecewise regressions using the ‘segmented’ R package (Muggeo, 2008).

## Results

### Bone properties with age

Mean trabecular properties calculated in the whole calcaneus are plotted against age in Figure 2. At birth, BV/TV is relatively high followed by a sharp decline in BV/TV during the first month. BV/TV gradually begins to increase between the first and second months for roughly two years after which BV/TV flattens out. Tb.Th and Tb.Sp increase gradually with age from birth to seven years of age (the maximum age of juveniles in the sample). There is no relationship between age and whole bone average degree of anisotropy. Variation in all variables is greater between older individuals.

**Figure 2.**
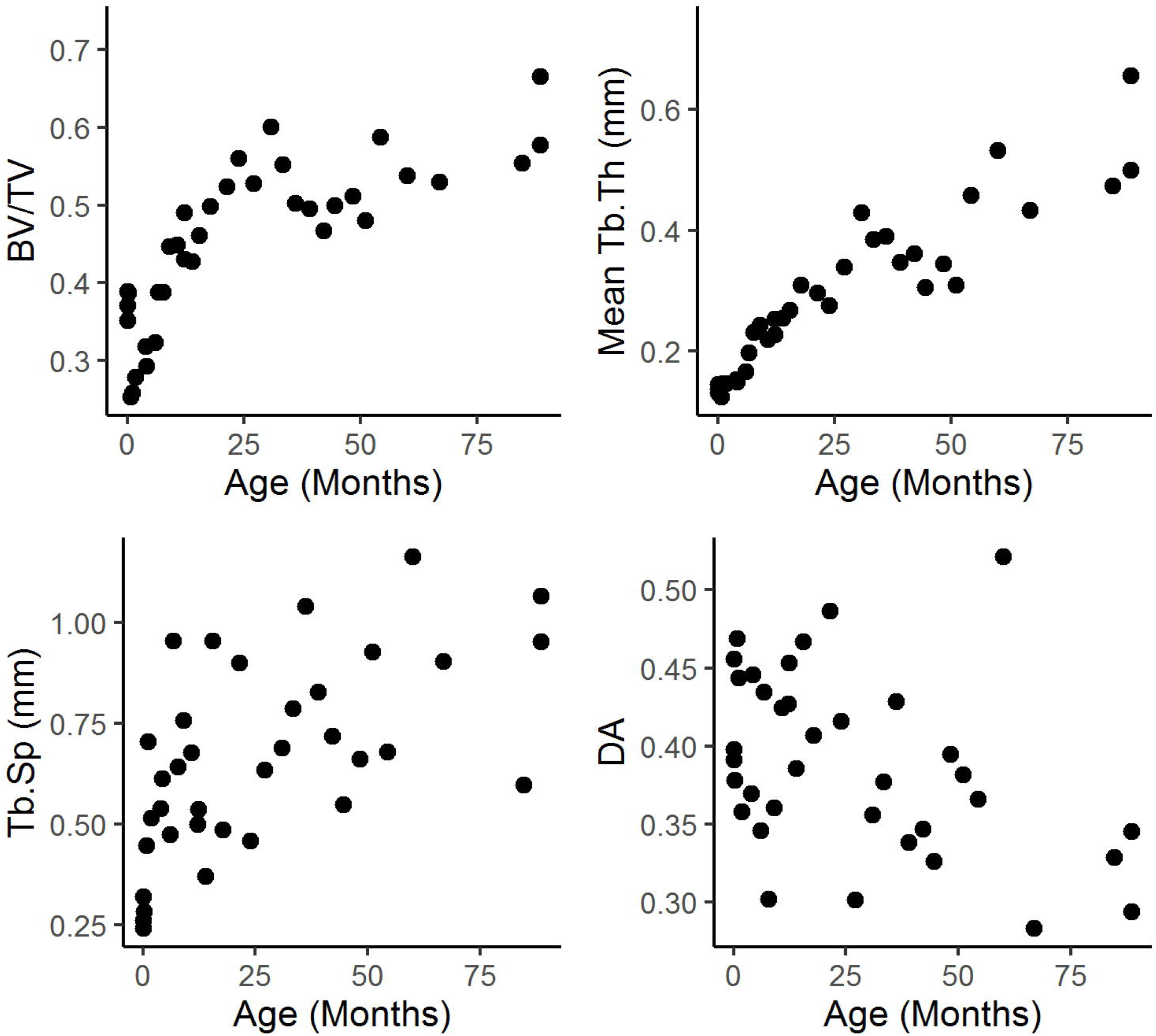
Whole bone trabecular properties plotted against age in months. BV/TV = bone volume fraction, Tb.Th = trabecular thickness, Tb.Sp = trabecular spacing, DA = degree of anisotropy.

Trends can be clearly observed when the data are plotted on the log-log scale (Figure 3). The drop in BV/TV is correlated with an increased average distance between trabeculae (Tb.Sp) while mean trabecular thickness remains constant. After the first month of life, when macaques start to locomote somewhat regularly, BV/TV rises in concert with Tb.Th and keeping a relatively constant Tb.Sp. Indeed, change in BV/TV is very strongly correlated to Tb.Th (linear regression R^2^=0.85, p<2.2e-16, AIC=-128.7). Adding an interaction between Tb.Th and Tb.Sp increases R^2^ to 0.88 and has a slightly lower AIC of −133.8, indicating a higher model quality. These data suggest that much of the variation in BV/TV can be explained by an interaction between average trabecular thickness and average distance between trabeculae.

**Figure 3.**
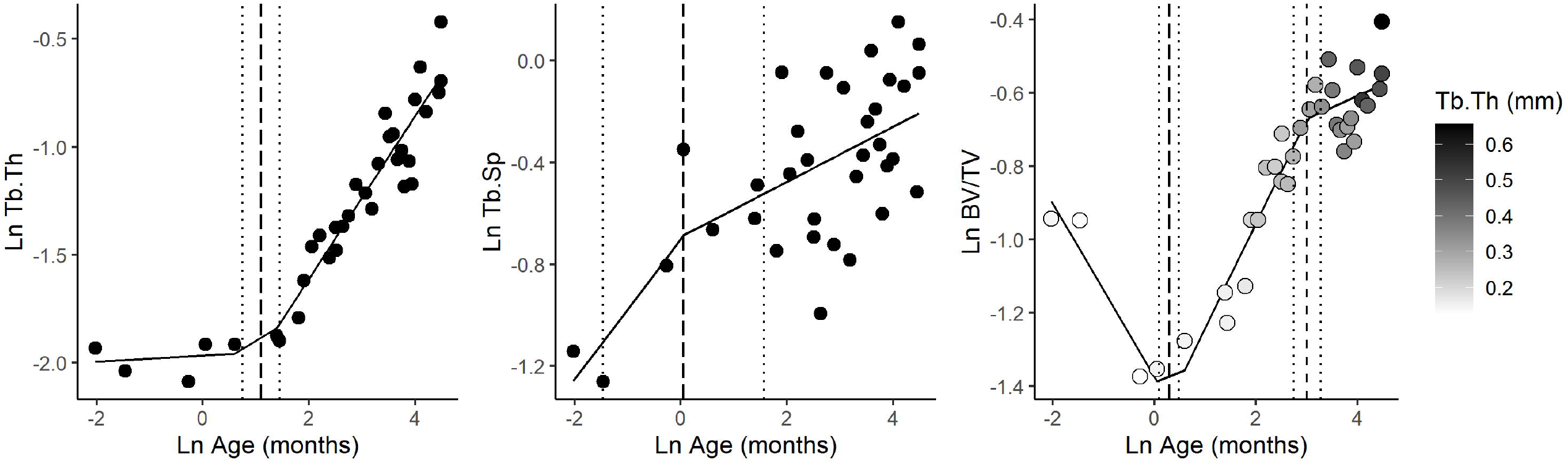
Plots showing the natural log of BV/TV, Tb.Th, and Tb.Sp plotted against the natural log of age. Piecewise regression output is visualized as solid lines. The dashed lines represent the maximum likelihood of the break points, and the dotted lines represent the standard error of the estimate. In the BV/TV plot points are filled based on trabecular thickness. BV/TV = bone volume fraction, Tb.Th = trabecular thickness, Tb.Sp = trabecular spacing.

We performed a piecewise regression model and used maximum likelihood to identify potential transition points in the data (Figure 3). The piecewise regression model (ln BV/TV ∼ ln age) has a higher quality (adj. R^2^=0.89, AIC=-65.9) compared to a regular linear regression model (R^2^=0.61, AIC=-24.6) Break points were identified at 1.3 months with a standard error of 1.2 months (ln 0.29, s.e. 0.19), and one at 20.2 months with a standard error of 1.2 months (ln 3.01, s.e. 0.27). The first break point coincides with the period of locomotor onset in Japanese macaques. The second break point overlaps with the time range where young macaque’s locomotor skill and repertoire approaches that of adults. For Tb.Th, (ln Tb.Th ∼ ln age) a model with a single break fits best (adj. R^2^=0.92, lowest AIC=-36), and fits better than the standard regression model (R^2^=0.81, AIC=-10.9). Here the breakpoint is identified at 2.9 months with a standard error of 1.4 months (ln 1.09, s.e. 0.35). While slightly later than the first break point identified in BV/TV, they overlap in standard error. For Tb.Sp, (ln Tb.Sp, ln age) a piecewise regression model has a lower R^2^ and a higher AIC than the linear regression, although the two models are virtually equal. The breakpoint occurs at 1.1 months with a standard error of 4.5 months (ln 0.05, s.e. 4.53).

### Bone properties with body weight

Log-log plots of trabecular properties against body weight at death are shown in Figure 4. Regression coefficients for a linear model with body weight as a predictor and trabecular properties as dependent variables are provided in Table 2. Significant positive relationships were found between body weight and BV/TV, Tb.Sp, and Tb.Th. For Tb.Th and Tb.Sp the predicted isometric scaling coefficient is 1/3 as body mass scales volumetrically (to the third power) and trabecular thickness scales by length (to the first power). As DA and BV/TV are both ratios they should scale with a slope of 0 under isometry. BV/TV and Tb.Th scale with positive allometry (Table 2) while Tb.Sp scales with slight negative allometry.

**Figure 4.**
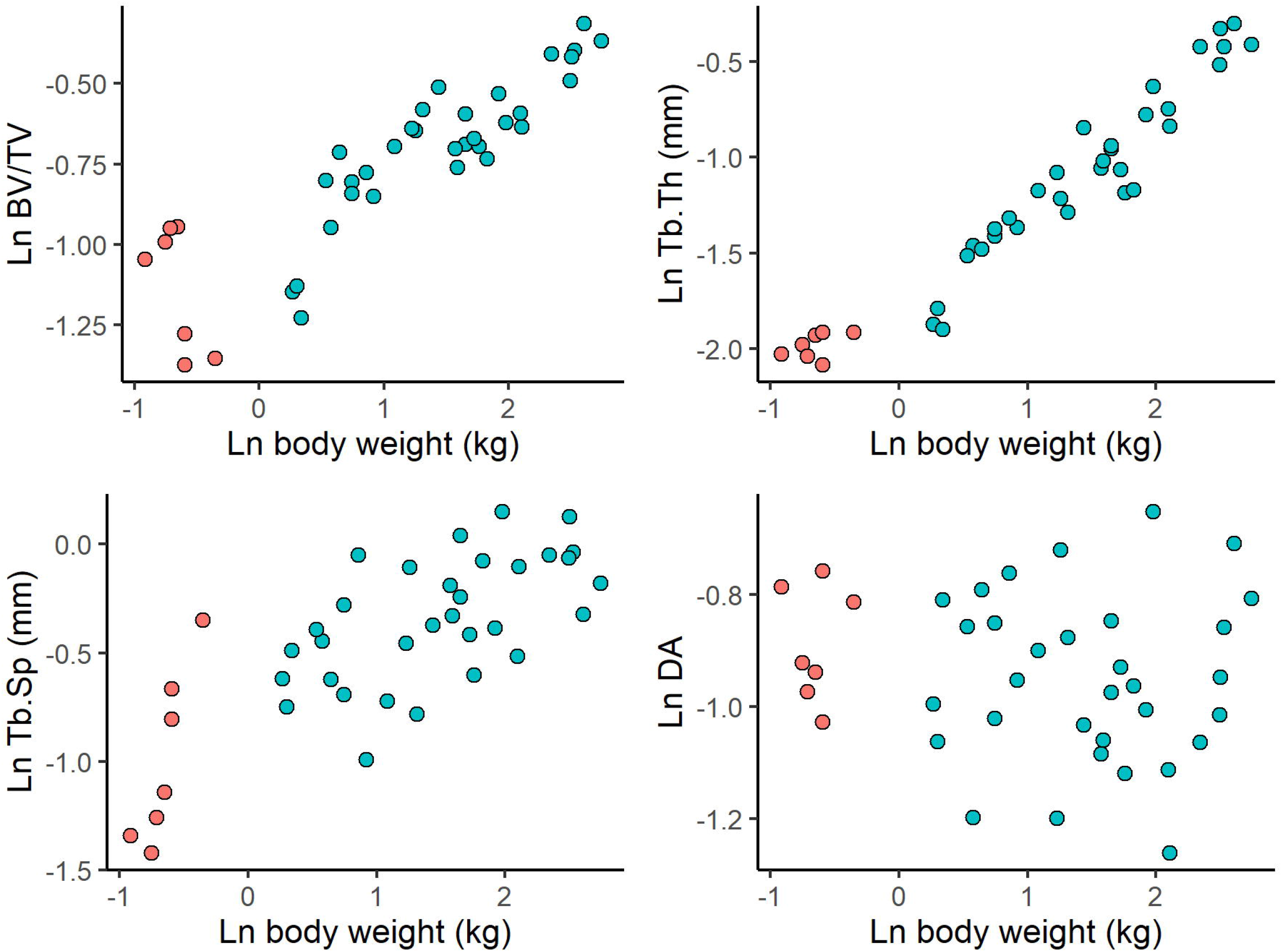
Natural log of body weight at death plotted against the natural log of trabecular properties. Red dots are younger than 2 months and blue dots represent individuals older than 2 months. BV/TV = bone volume fraction, Tb.Th = trabecular thickness, Tb.Sp = trabecular spacing, DA = degree of anisotropy.

**Table 2.** Regression coefficients of ln trabecular properties as dependent variable and ln body mass as the predictor.

| Dependent variable | Intercept | Slope | Slope s.e. | df | *p* | R<sup>2</sup> | AIC | Allometry |
| --- | --- | --- | --- | --- | --- | --- | --- | --- |
| **Ln BV/TV** | -1.012 | 0.225 | 0.020 | 37 | <0.001 | 0.77 | -42.65 | Positive |
| **Ln Tb.Th** | -1.756 | 0.487 | 0.022 | 37 | <0.001 | 0.93 | -37.59 | Positive |
| **Ln Tb.Sp** | -0.770 | 0.284 | 0.039 | 37 | <0.001 | 0.58 | 9.47 | Negative |
| **Ln DA** | -0.921 | -0.017 | 0.022 | 37 | 0.449 | 0.02 | -36.28 | None |

BV/TV = bone volume/total volume, Tb.Th = average trabecular thickness (mm), Tb.Sp = average distance between trabeculae (mm), DA = degree of anisotropy.

Regression parameters for models where trabecular properties are predicted by the interaction between body weight and locomotor independence are given in Table 3. These interaction models generate a slope and intercept for specimens that younger than two months who are fully or partially dependent on their mothers for locomotion, and those older than 2 months who are generally independent.

**Table 3.** Regression coefficients of ln trabecular properties as dependent variable to an interaction between body weight and locomotor independence predictors. Sample size is 39 for all models.

| Dependent variable | Locomotor independence | Intercept | Slope | *p* | R<sup>2</sup> | AIC |
| --- | --- | --- | --- | --- | --- | --- |
| **Ln BV/TV** | Dependent | -1.635* | -0.764* | <0.001 | 0.826 | -52.696 |
|  | Independent | -1.067* | 0.259* |  |  |  |
| **Ln Tb.Th** | Dependent | -1.862 | 0.186 | <0.001 | 0.947 | -46.518 |
|  | Independent | -1.901* | 0.571* |  |  |  |
| **Ln Tb.Sp** | Dependent | 0.376* | 2.094* | <0.001 | 0.664 | 1.770 |
|  | Independent | -0.679* | 0.229* |  |  |  |
| **Ln DA** | Dependent | -0.875 | 0.019 | 0.790 | 0.000 | -32.816 |
|  | Independent | -0.954 | 0.002 |  |  |  |

\*Coefficient significantly different from 0. BV/TV = bone volume/total volume, Tb.Th = average trabecular thickness (mm), Tb.Sp = average distance between trabeculae (mm), DA = degree of anisotropy.

The linear models with an interaction between body mass and locomotor independence given in Table 3 have higher R^2^ and lower AIC for BV/TV, Tb.Th, and Tb.Sp compared to the linear models presented in Table 2. This indicates that the interaction models have a greater fit to the data while at the same time not being overfit compared to the univariate models. Tb.Th is predicted by body mass only after locomotor independence. BV/TV scales with a negative slope before locomotor independence and with a positive slope afterwards. Tb.Sp scales with a positive slope before and after locomotor independence but the slope is significantly shallower after locomotor independence.

### Morphometric maps

The trabecular properties presented above were averaged across the whole calcaneus. Figure 5 shows the heterogeneous distribution of BV/TV and primary Young’s modulus (primary direction of bone stiffness) in sagittal cross sections of a subset of specimens. When BV/TV is scaled between the sample minimum of 0.14 and maximum of 0.70, trabecular structure appears relatively homogeneous at birth and becomes increasingly heterogeneous with age. When the colormaps are scaled to the minimum and maximum values for each individual, regional variation is evident at all stages. BV/TV is greatest under the joint surfaces, particularly of the posterior talar facet.

**Figure 5.**
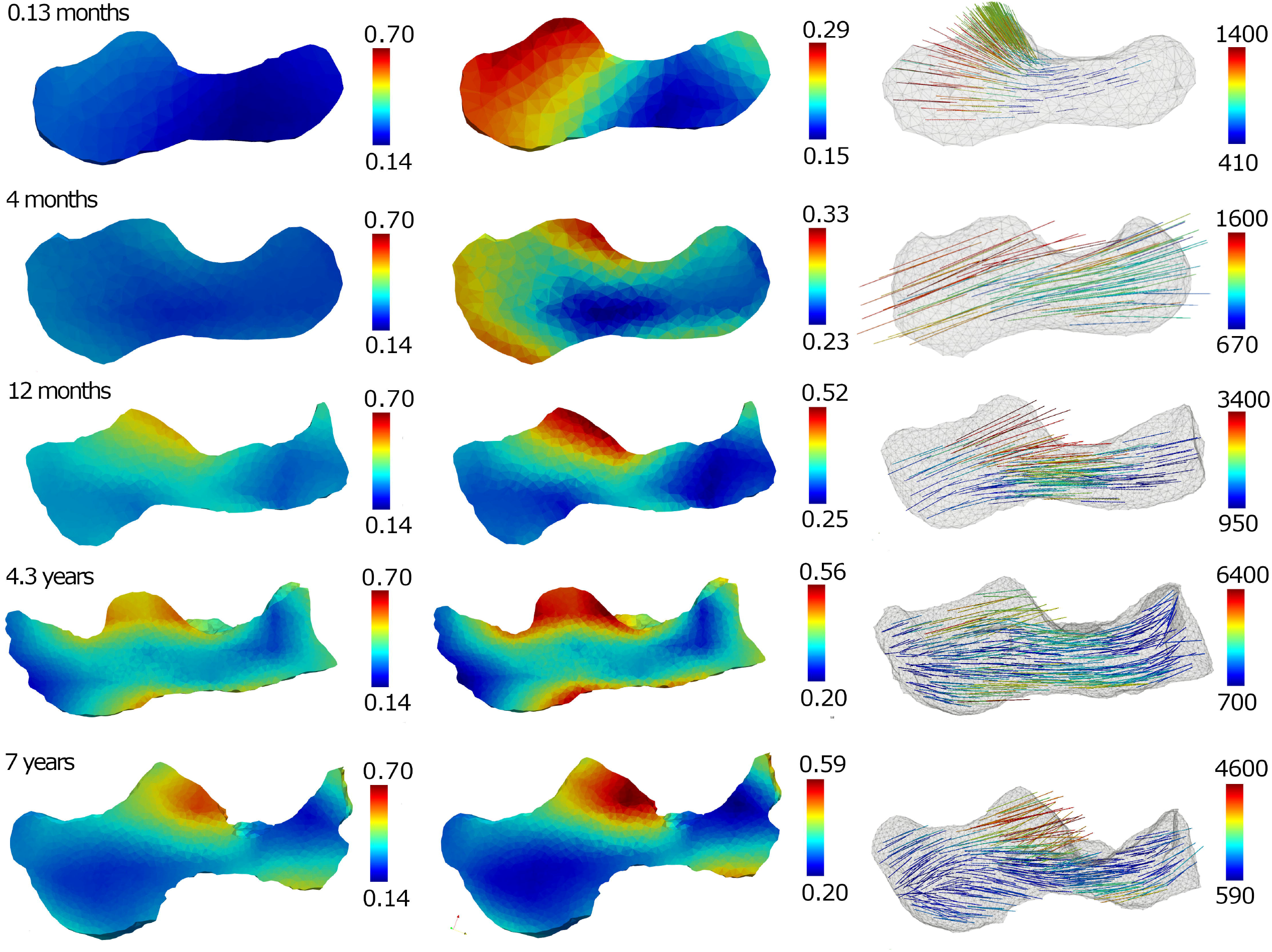
BV/TV scaled between the sample minimum of 0.14 and maximum of 0.70 (left). BV/TV scaled to the minimum and maximum of each individual calcaneus (center). Vectors show primary direction and magnitude of Young’s modulus (MPa) for each individual (right). Bones are not to scale.

The primary Young’s modulus represents the direction of maximum bone stiffness taken at points uniformly distributed throughout the calcaneus (Figure 5 and 6). At birth, primary Young’s modulus is in the direction in which the bone is growing, orthogonal to the growth plate (Figure 6, bottom left). However, the primary direction of bone stiffness differs in individuals past the first month of age, when the macaques start to locomote regularly (Figure 6, top left). Measured from 12 o’ clock counterclockwise in a sagittal slice through the calcaneus underneath the posterior talar facet, the direction of the primary Young’s modulus changes from 15-35 degrees to 100-110 degrees in all individuals aged greater than 1.8 months of age (Figure 6, right).

**Figure 6.**
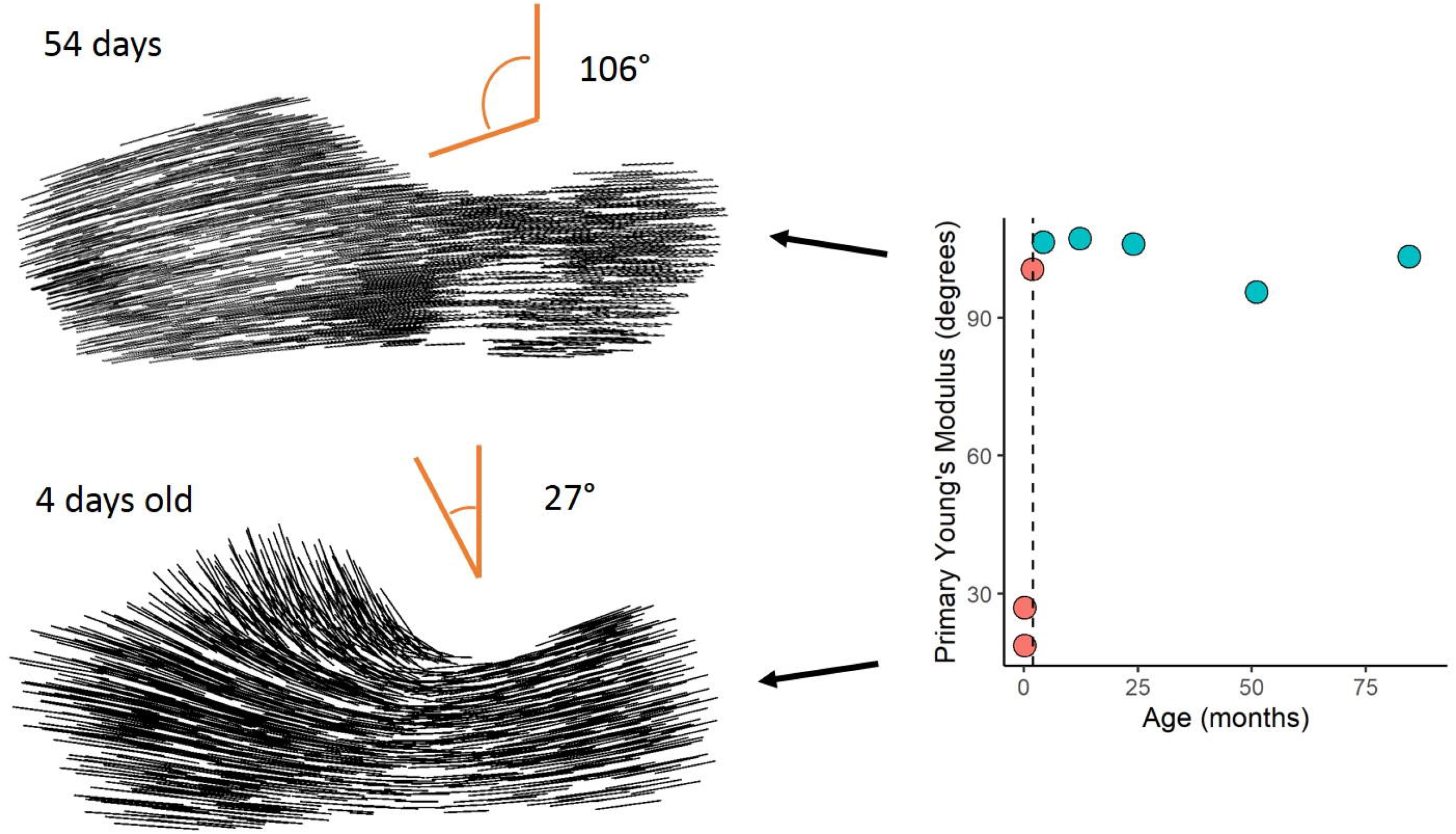
Sagittal slices through the calcaneus with black lines representing the direction in which the Young’s modulus is greatest. On the right the angle of the primary Young’s modulus is plotted against age (months) with the dotted line representing the timing of locomotor independence.

### Bones and brains

Figure 7 shows the close overlap between attainment of adult brain size and adult-like BV/TV. While macaques reach their maximum adult body size between 10 and 12 years of age (Hamada, 1994), 95% of maximum brain size is reached at two years of age and 100% after 5 years. BV/TV continues to increase slightly after the age when adult brain size has been obtained and body weight continues to increase.

**Figure 7.**
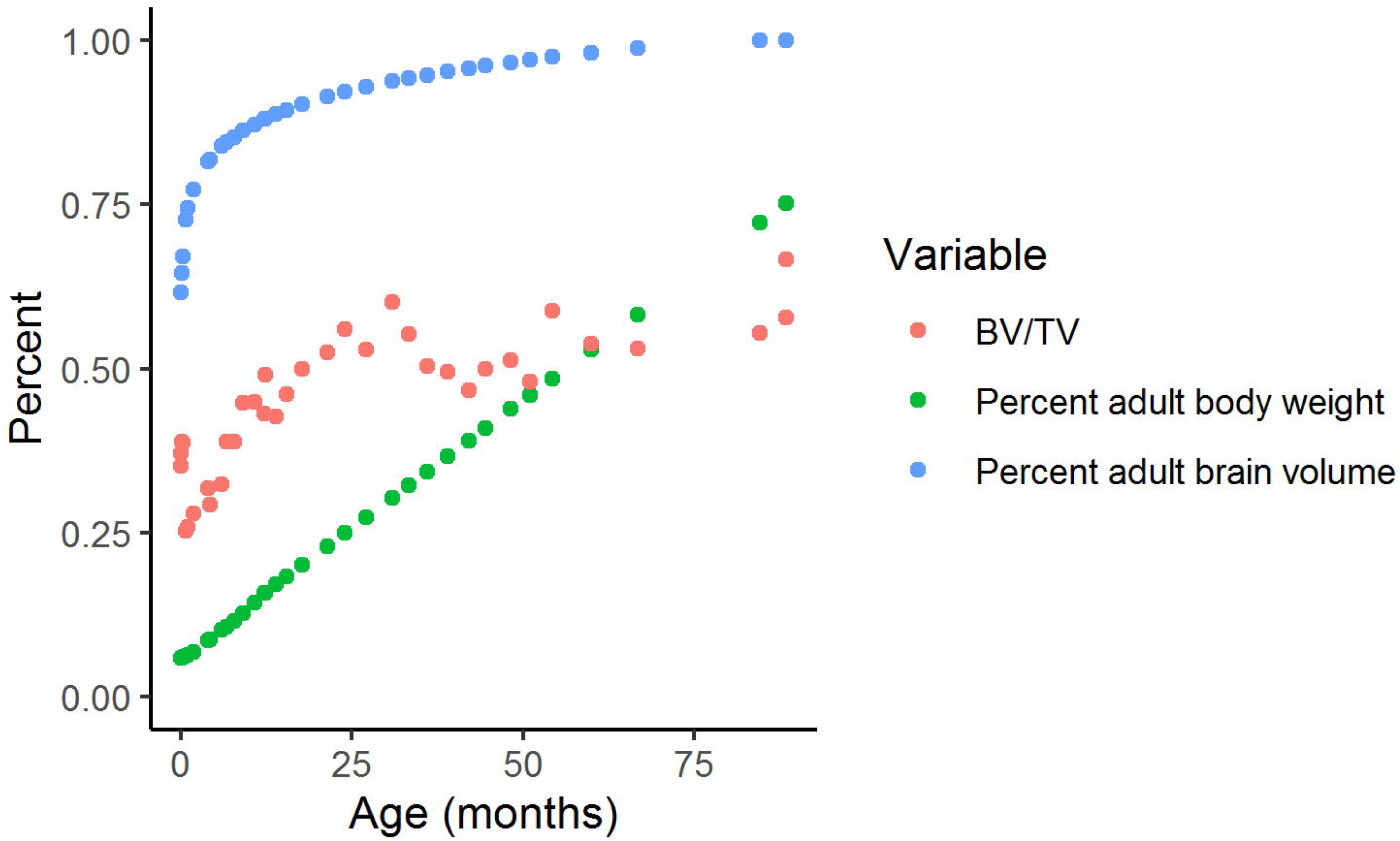
Plot of changes with age in calcaneus BV/TV, estimated percentage adult brain size, and estimated percentage of adult body weight.

If we assume that there is a relationship between brain development and locomotion, we can explain the patterns in Figure 7 with the following model: while the brain is still growing, increases in neuromuscular control and locomotor experience make loading environment of the calcaneus, and by proxy trabecular structure, increasingly similar to that of adults. When the brain has reached its full size, neuromuscular control of locomotion and locomotor loading conditions also approach the adult-like pattern (SOM Table S.1., SOM Figure S.1, S.2). This model suggests that the steep early age-related variation in BV/TV found in the macaque calcaneus may be related increasing neuromuscular control of gait with a slight positive allometry after gait has matured but when body mass continues to increase. We test this model using percent adult brain size as a proxy for neuromuscular control. In Table 4 we compare various types of models to assess which model predicts trabecular properties best (lowest AIC, highest R^2^).

**Table 4.** Comparison of various regression models for each trabecular property, the best fit (highest R^2^) and highest model quality (lowest AIC) are in bold. All continues variables are natural log transformed and n=34 for all models after removing missing cases due to unknown age.

| Dependent | Independent | Adj. $R^2$ | AIC | p |
| --- | --- | --- | --- | --- |
| BV/TV | BM | 0.69 | -32.9 | 1.08e-9 |
|  | BM * independence | 0.76 | -40.9 | 4.3e-10 |
|  | BM * percent adult brain size | 0.73 | -37.0 | 2.4e-9 |
|  | **BM \* percent adult brain size \* independence** | **0.85** | **-54.1** | **1.3e-10** |
| Tb.Th | BM | 0.90 | -33.7 | 2.2e-16 |
|  | **BM \* independence** | **0.92** | **-37.8** | **2.2e-16** |
|  | BM * percent adult brain size | 0.92 | -36.1 | 2.2e-16 |
|  | BM * percent adult brain size * independence | 0.92 | -32.5 | 6.5e-14 |
| Tb.Sp | BM | 0.54 | 11.4 | 6.7e-7 |
|  | **BM \* independence** | **0.63** | **4.9** | **2.9e-7** |
|  | BM * percent adult brain size | 0.59 | 8.7 | 1.4e-6 |
|  | BM * percent adult brain size * independence | 0.62 | 9.1 | 1.8e-5 |
| DA | BM | 0.09 | -32.8 | 0.09 |
|  | BM * independence | 0.00 | -29.0 | 0.40 |
|  | BM * percent adult brain size | 0.01 | -29.3 | 0.35 |
|  | BM * percent adult brain size * independence | 0.00 | -23.7 | 0.65 |

Abbreviations: BM = body mass at death (kg), independence = older than 2 months, independent if older than two months, BV/TV = bone volume/total volume, Tb.Th = average trabecular thickness (mm), Tb.Sp = average distance between trabeculae (mm), DA = degree of anisotropy.

Variation in BV/TV is best explained by a 3-way interaction between body mass, percentage of adult brain size, and locomotor independence. Trabecular thickness and spacing are best explained by a 2-way interaction between body mass and locomotor independence. Degree of anisotropy is not correlated with any variables tested above; this is not surprising since DA reflects more local changes in trabecular structure that likely get averaged out when analyzed in the whole calcaneus.

## Discussion

We made the following predictions:

1. **Onset of locomotion**: we predicted that the slope of BV/TV with age in our sample should shift from negative to positive, that trabeculae would become thicker on average, and that the Young’s modulus should change direction from orthogonal to the ossification centre to joint surfaces. *All of these predictions were confirmed*.
2. **Appearance of adult-like locomotor repertoire:** When an adult-like locomotor repertoire is achieved between the ages of 1 and 4, the predicted positive slope in BV/TV was predicted to follow a shallower slope due to allometric scaling with still increasing body size. *This prediction was confirmed*.
3. **Neuromuscular maturation of gait:** We predicted that trabecular properties at different ages should be predictable by an interaction between body mass and percent adult brain size. *This prediction was confirmed in BV/TV after adding in an extra interaction term for locomotor independence. The prediction was not confirmed for average trabecular thickness or spacing*.

### Results in the context of the wider literature

The developmental trajectories of trabecular properties in the calcaneus of Japanese macaques resemble those of other mammals (Wolschrijn and Weijs, 2004; Tsegai et al., 2018; Colombo et al., 2019; Ragni, 2020) including humans (Ryan and Krovitz, 2006; Gosman, 2007; Raichlen et al., 2015; Ryan et al., 2017; Saers et al., 2020), indicating a generally shared mechanism of growth (Carter and Beaupré, 2001). The distribution of BV/TV is substantially more homogenous in younger individuals and becomes increasingly heterogeneous with age, similar to results reported by Tsegai et al. (2018) for the chimpanzee postcranium. The development of trabecular structure of the calcaneus of Japanese macaques follows the same patterns as the human calcaneus, but with differences in the timing of stages (Saers et al., 2020), as well as other postcranial elements (Ryan and Krovitz, 2006; Gosman and Ketcham, 2009; Raichlen et al., 2015; Saers et al., 2020). In both species bone is overproduced during early development with high BV/TV and struts largely oriented perpendicular to the ossification center in the calcaneus, or the growth plate in long bones. In both species BV/TV reduces after birth and begins to increase again at the same time when individuals typically begin independent locomotion. At the same time the primary Young’s modulus shifts in direction from orthogonal to the growth plate to joint surfaces. This reorientation in the direction in which the bone is loaded helps to more efficiently distribute the loads placed upon the calcaneus during locomotion (Wolff, 1867; Roux, 1881; Zysset, 2003; Maquer et al., 2015). Contrary to our findings, Tsegai et al. (2018) did not find the initial overproduction of trabecular bone, followed by a drop in BV/TV. However, their sample included just one individual that was possibly younger than 6 months, potentially causing them to miss drop in BV/TV shortly after birth that has been reported in humans and now macaques.

### The link between bones and brains

Locomotor patterns are transformed from an immature state to increasingly adult-like patterns during development. Changes in loading patterns with advancing age are generated by neural circuits that develop in parallel to increases in physical size and weight of a growing organism. Our results lend strong, but indirect support, for a model where trabecular structure is the product of age-related variation in loading conditions. These changes in loading conditions are a product of the development of gait which, in turn, is the product of neural maturation with age and experience. If this link is demonstrated experimentally, and in other species, trabecular structure could be used as a proxy not for just development of locomotion, but also neural maturation in fossil species. This brain-bone connection would then serve as a powerful life history marker in fossil species.

While the biomechanical data required to test the association between percent adult brain size and locomotor loading in macaques does not exist in the literature, we can test the association with human data. If our model of trabecular bone adaptation is correct, there should be a close relationship between neuromuscular control of gait, gait mechanics, and underlying trabecular structure in humans. We briefly tested this association using experimental data from other publications (Vaughan et al., 2003; Müller et al., 2012, see SOM S.1). In human children aged 1-13 years old, experimentally measured force-time-integral of normal walking (Müller et al., 2012), normalized by body mass, is perfectly correlated with an experimentally derived measure of neuromaturation (Vaughan et al., 2003) (R^2^=0.99, see SOM Table S.1). This measure of neuromaturation again is correlated almost perfectly linearly with estimated percentage of adult brain size in humans (R^2^=.98, percent brain size data calculated from (Cofran and Desilva, 2015)). These results suggest an extremely tight correspondence between the mechanics of foot loading relative to body size, locomotor development, neuromaturation, and estimated percentage of adult brain size. These perfect correlations are especially striking because they arise between three different measures derived from different experiments in different papers and on divergent topics (Vaughan et al., 2003; Müller et al., 2012; Cofran and Desilva, 2015). However, future experimental work is required to causally link these strong correlations.

### Using trabecular ontogeny as a life history marker

While some aspects of human bone morphology are genetically determined, others are environmentally induced. For example, the human lateral patellar lip is present already at birth (Scheuer and Black, 2004; Lovejoy, 2007), whereas the human bicondylar angle develops postnatally in response to mechanical loading associated with bipedal locomotion (Tardieu and Trinkaus, 1994; Tardieu, 1999, 2010). In terms of trabecular bone, Barak et al. (2011) showed experimentally that differences in peak loading angle as well as magnitude alter trabecular bone orientation and BV/TV in sheep. Our ontogenetic data also suggest that age-related variation in trabecular structure corresponds to variation in loading conditions during landmark events in the maturation of gait (onset of locomotion, adult-like gait). As such, our results suggest that trabecular structure can potentially be used to infer the timing of locomotor onset and the achievement of adult-like locomotor repertoires. The advent of independent locomotion coincides with important aspects of mammalian brain size and neuromuscular development (Campos et al., 2000; Garwicz et al., 2009; Anderson et al., 2013). Patterns of locomotor development may therefore provide unique insights into the evolution of locomotor modes, how locomotion develops, neuromaturation, the onset of independent locomotion in fossil species (Figure 8).

**Figure 8.**
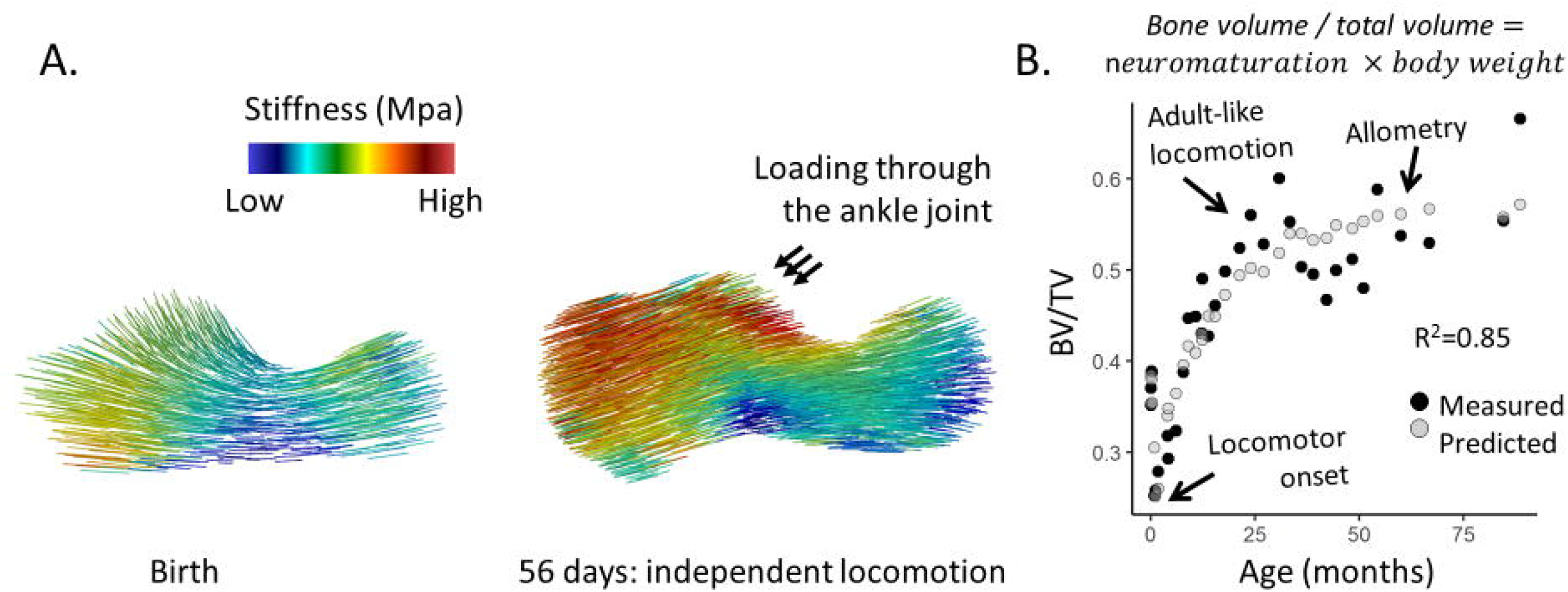
Summary of the paper findings. (Left) Material stiffness direction and magnitude at birth and at locomotor independence. (right) BV/TV plotted against age in months. The black dots represent measured BV/TV, gray dots are predicted BV/TV based on the 3-way interaction between locomotor independence, percent adult brain size, and body mass.

### Limitations

The data presented here correspond to a model where brain maturation increases neuromuscular control of gait, which, in turn, affects the mechanics of gait, which then shape loading patterns of the foot to which trabeculae dynamically adapt. However, our study is cross-sectional with a skeletal sample of individuals of whom we do not know the individual behavior during life. As such we cannot quantify the changes in mechanical loading over time directly. This study design cannot be used as evidence of a causal mechanism. Confirming our proposed model will require controlled experiments resulting in longitudinal data, ideally in several species.

The percentage of adult brain size is a very rough proxy for neural development and numerous changes in brain composition and wiring occur after adult brain size has been reached (Lebel and Beaulieu, 2011). However, as shown in the appendix, percent adult brain size in humans (calculated from Cofran and Desilva, 2015) is perfectly correlated (R^2^=0.99) with experimentally derived measures of neuromaturation (Vaughan et al., 2003). This simple measure of percentage adult brain size is all that is available in paleontological contexts and it is therefore encouraging to that we can report such tight correlations.

## Conclusions

The developmental trajectories of trabecular properties in the calcaneus of Japanese macaques are similar to other species, indicating a broadly shared mechanism of growth. Trabeculae are overproduced at birth, followed by refinement leading to adaptation to local conditions and resulting in a species and joint specific heterogeneous state. After the first month of age, when the macaques begin to regularly locomote coincides with striking changes in trabecular structure, including a reorientation of the primary Young’s modulus from orthogonal to the growth plate to the direction of joint surfaces.

The results indicate that variation in trabecular morphology closely corresponds to the adoption of independent locomotion in Japanese macaques. Locomotor independence coincides with important aspects of mammalian brain size and neuromuscular development. Such bony markers of locomotor development could therefore be compared with other developmental milestones that track the overall pace of life history in fossil species. If the model presented in this paper holds up under longitudinal experimental conditions, trabecular structure can be used both to infer behavior from fossil morphology (Barak et al., 2013; Tsegai et al., 2013; Skinner et al., 2015; Kivell, 2016; Stephens et al., 2016; Bishop et al., 2018; Ryan et al., 2018; Sorrentino et al., 2019; Dunmore et al., 2020) and to serve as a valuable proxy for neuromuscular maturation and life history events like locomotor onset and the achievement of an adult-like gait.

## Acknowledgements

Many thanks to Dr. Takeshi Nishimura for providing access to the skeletal collection, databases, and CT scanner at the Primate Research Institute, Inuyama, Japan.

## Data availability statement

The data that support the findings of this study are available on request from the corresponding author.

## Author contributions

JPPS: concept/design, acquisition of data, data analysis/interpretation, drafting of the manuscript.

ADG: critical revision of the manuscript and statistical advice. TMR, JTS: funding acquisition and critical revision of the manuscript.

## Supplementary material

### Relationship between gait mechanics and neuromaturation in humans

The human brain reaches 95% of its adult size around the age of six (Fair and Schlaggar, 2008). This timing coincides with human gait becoming almost exactly like adults, with some differences in gait based on allometric differences in body proportions between adults and juveniles (Sutherland, 1997). During the ontogeny of locomotion two parallel forms of growth occur: the infants physical size (height and weight) increases; and neural circuity develops which transforms the infants locomotor pattern (Forssberg, 1985). During the early ontogeny of gait, infants are mainly focused on minimizing the risk of falling. When individuals increase in strength stability improves and postural constraints are reduced (Vaughan et al., 2003). It is thought that development subsequently proceeds to select the most optimal neural networks (Forssberg, 1999), resulting in a reduction in the variability of muscle activation and co-contraction, and the adult gait pattern emerges (Okamoto et al., 2003).

An equation to estimate neuromaturation of human locomotion was derived by Vaughan et al. (2003). They show that non-dimensional scaling of velocity accounts for physical growth, yielding insight into the process of neuromaturation. The equation works by estimating dimensionless velocity (β) from dimensionless step frequency and dimensionless step length. This dimensionless approach removes body dimensions from the equation, what remains is the signal of neuromuscular maturation. Neuromaturation of locomotion for age was calculated with an equation derived by Vaughan et al. (neuromaturation (β) = 0.45 * (1-e^-0.05*t^) where t = age in months). Encouragingly, this derived measurement of neuromaturation is correlated almost perfectly with our estimate of percentage of adult brain size (R^2^=.98, p=1.45e-10).

According to our model, BV/TV of the calcaneus should be predictable largely by its loading environment (Carter et al., 1996). This loading environment can be roughly quantified by force-time-integrals and peak plantar pressure data. Müller et al. (2012) measured force-time intervals (FTI, newtons*second, the area under a force-over-time curve) in children aged between 1 and 13 years. Our model also predicts that the loading environment should vary depending on neuromuscular control as juveniles learn to walk, and allometric scaling of body mass and limb proportions. To test whether this model is accurate we combine data from two different studies; neuromaturation for age from Vaughan et al. (2003) and force-time-integrals and peak plantar pressure for age from Muller et al. 2012) for children aged between 1 and 13 years old. We estimated percentage of adult brain size using our equation derived from data by Cofran and DeSilva (2015). We test whether peak plantar pressure and body mass normalized force-time-integrals can be predicted by body mass, neuromaturation, or an interaction between body mass and neuromaturation (or percent adult brain size). In all cases the interaction between neuromaturation and body mass perfectly predicts both peak plantar pressure and body mass normalized force-time-integrals in both the hindfoot and the whole foot (SOM Table S.1).

### Peak plantar pressure

Predictions of FTI/BM and peak plantar pressure data from the models are plotted in SOM Figure S.1. Based on these very strong correlations we infer that the interaction between neuromaturation (and by proxy, percent adult brain size) and body mass accurately predicts the force-time-integral (loading environment) of the foot, both peak plantar pressure in the whole foot and hindfoot, and body mass normalized force-time-integrals in the whole foot and hindfoot. These plots support our model where the loading environment of the foot can be accurately predicted by the interaction between neuromaturation (or by proxy, percent adult brain size) and body mass.

The force-time integral relative to body size (FTI/BM) with age can be almost perfectly predicted by the level of neural maturation (or percent brain size) and body size of an individual. This very strongly supports our model that neural maturation could be an important factor that contributes to changing loading patterns as gait develops. Furthermore, BV/TV in the human calcaneus [taken from Saers et al. (2020)] is predicted by the same interaction between body mass and neuromaturation (Adj. R^2^=.85, p= 2.7e-5, see Figure S2). At least some of the error in BV/TV underlying the lower R^2^ can be explained by the fact that BV/TV was measured in a small sample of archaeological specimens for which age and body size had to be estimated. Meanwhile, FTI and neuromaturation were calculated from equations based on mean trends in large samples from the literature, and therefore do not contain variation that would be associated with intrinsic individual differences and variation in the tempo at which individuals mature, which is there in BV/TV. Overall, this correlational evidence is highly encouraging and future experimental longitudinal study may be able to show the exact causal mechanism.

## Tables

**SOM Table S.1.** Comparison of various regression models predicting loading conditions of the foot of 1-13-year-old children. Neuromaturation and percent adult brain size estimated based on age.

| Dependent | Independent | Adj. $R^2$ | AIC | p |
| --- | --- | --- | --- | --- |
| FTI/BM<br>hindfoot | BM | 0.71 | -63.4 | 3.1e-4 |
|  | Neuromaturation | 0.90 | -77.4 | 7.4e-7 |
|  | **BM \* Neuromaturation** | **0.99** | **-110.8** | **1.8e-10** |
|  | BM * percent adult brain size | 0.98 | -99.6 | 9.0e-9 |
| FTI/BM<br>whole foot | BM | 0.87 | -50.9 | 3.3e-6 |
|  | Neuromaturation | 0.75 | -42.4 | 1.3e-4 |
|  | **BM \* Neuromaturation** | **1.00** | **-97.3** | **6.9e-12** |
|  | BM * percent adult brain size | 0.99 | -81.2 | 1.8e-9 |
| Peak plantar pressure<br>hindfoot | BM | 0.69 | 125.0 | 4.0e-4 |
|  | Neuromaturation | 0.93 | 106.3 | 1.3e-7 |
|  | BM * Neuromaturation | 0.98 | 89.8 | 1.6e-8 |
|  | **BM \* percent adult brain size** | **0.98** | **89.4** | **1.4e-8** |
| Peak plantar pressure<br>whole foot | BM | 0.96 | 109.9 | 4.0e-9 |
|  | Neuromaturation | 0.64 | 139.0 | 1.1e-3 |
|  | **BM \* Neuromaturation** | **0.99** | **85.6** | **6.4e-11** |
|  | BM * percent adult brain size | 0.98 | 89.4 | 1.4e-8 |
| Peak plantar pressure<br>Hindfoot / BM | BM | 0.94 | -23.1 | 4.7e-8 |
|  | Neuromaturation | 0.39 | 6.9 | 0.22 |
|  | **BM \* Neuromaturation** | **0.99** | **-54.0** | **4.7e-11** |
|  | BM * percent adult brain size | 0.98 | -38.9 | 8.7e-9 |
| Peak plantar pressure<br>whole foot / BM | BM | 0.92 | -21.6 | 1.6e-7 |
|  | Neuromaturation | 0.55 | 1.7 | 3.7e-3 |
|  | **BM \* Neuromaturation** | **0.99** | **-51.3** | **1.9e-10** |
|  | BM * percent adult brain size | 0.99 | -47.9 | 6.2e-10 |

FTI/BM = force-time-integral normalized by body mass, BM = body mass (Newtons).

## Figure legends

**SOM Figure S.1.**
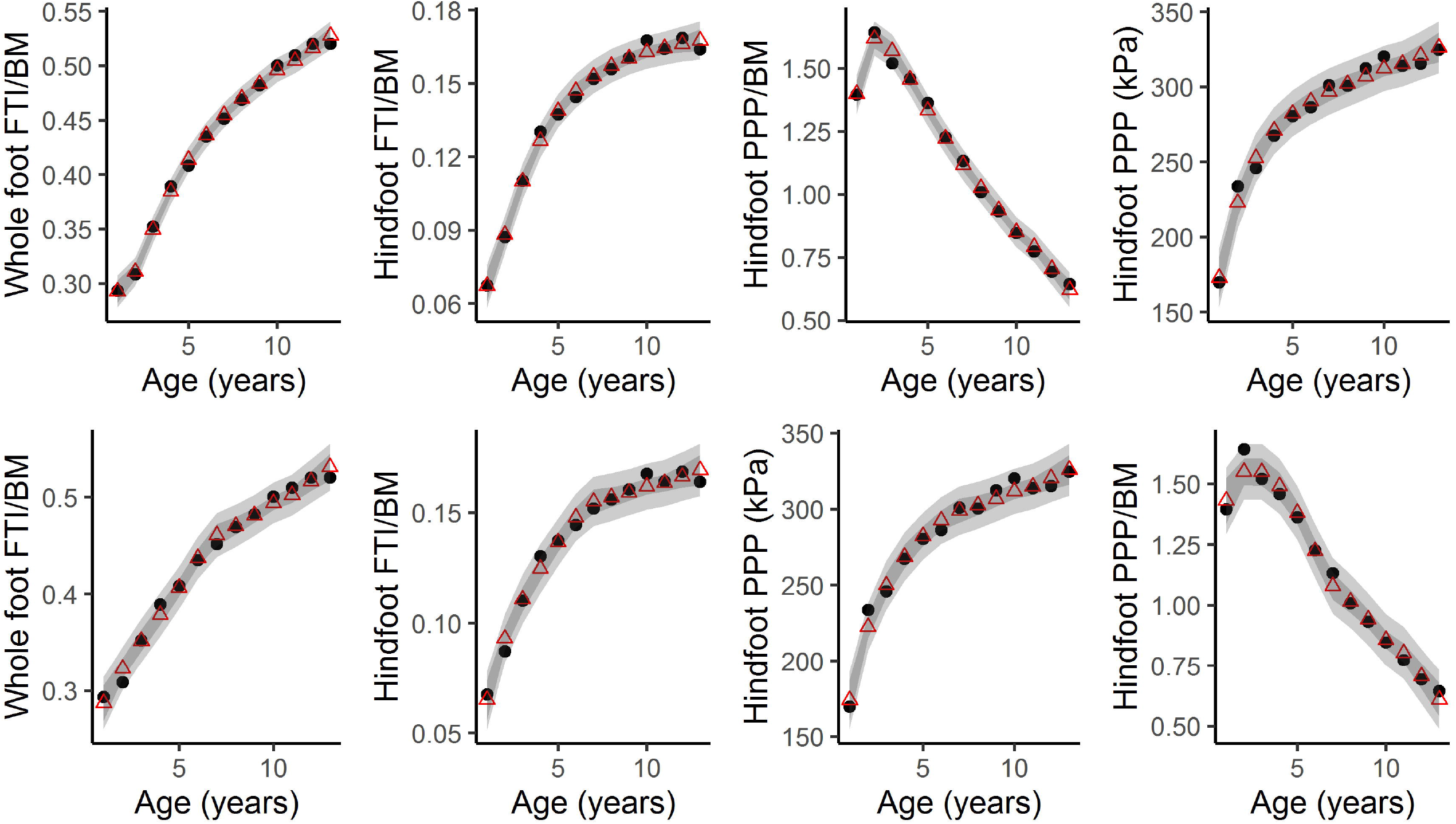
(Top) foot loading variables with age predicted by interaction between body mass and neuromaturation. (Bottom) foot loading variables predicted by interaction between body mass and percentage of adult brain size.

**SOM Figure S.2.**
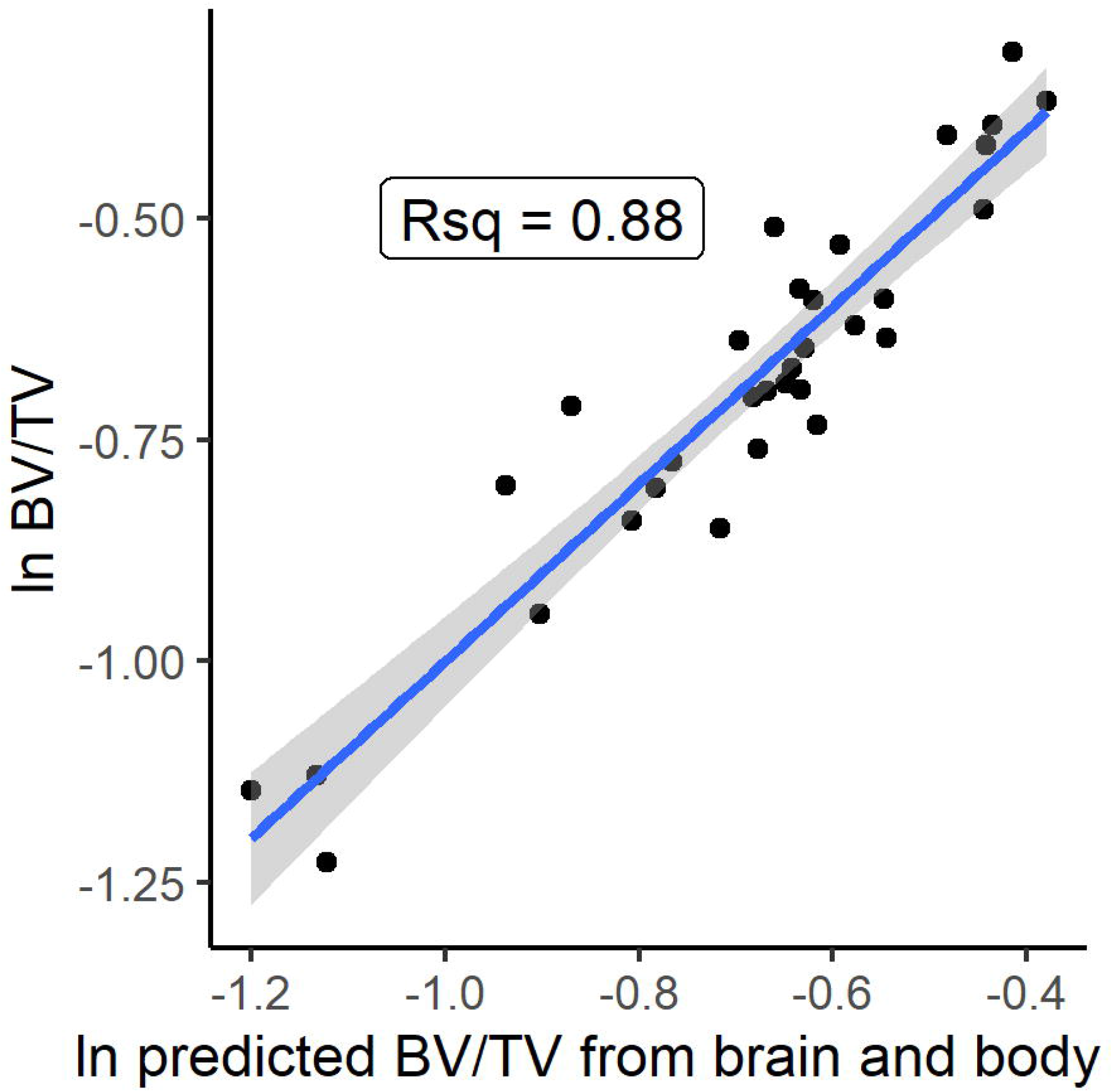
Regression between measured ln BV/TV (from Saers et al. (2020)) and BV/TV predicted by an interaction between neuromaturation and body size.

