## Supplementary data for "Growth and development of trabecular structure in the calcaneus of Japanese macaques (*Macaca fuscata*) reflects locomotor behavior, life-history, and neuromuscular development"

| **Dependent** | **Independent** | **Adj. $R^2$** | **AIC** | **p** |
|---|---|---|---|---|
| **FTI/BM** | BM | 0.71 | -63.4 | 3.1e-4 |
| **hindfoot** | Neuromaturation | 0.90 | -77.4 | 7.4e-7 |
| | **BM * Neuromaturation** | **0.99** | **-110.8** | **1.8e-10** |
| | BM * percent adult brain size | 0.98 | -99.6 | 9.0e-9 |
| **FTI/BM** | BM | 0.87 | -50.9 | 3.3e-6 |
| **whole foot** | Neuromaturation | 0.75 | -42.4 | 1.3e-4 |
| | **BM * Neuromaturation** | **1.00** | **-97.3** | **6.9e-12** |

| | | | | |
|---|---|---|---|---|
| | BM * percent adult brain size | 0.99 | -81.2 | 1.8e-9 |
| **Peak plantar pressure** | BM | 0.69 | 125.0 | 4.0e-4 |
| **hindfoot** | Neuromaturation | 0.93 | 106.3 | 1.3e-7 |
| | BM * Neuromaturation | 0.98 | 89.8 | 1.6e-8 |
| | **BM * percent adult brain size** | **0.98** | **89.4** | **1.4e-8** |
| **Peak plantar pressure** | BM | 0.96 | 109.9 | 4.0e-9 |
| **whole foot** | Neuromaturation | 0.64 | 139.0 | 1.1e-3 |
| | **BM * Neuromaturation** | **0.99** | **85.6** | **6.4e-11** |
| | BM * percent adult brain size | 0.98 | 89.4 | 1.4e-8 |
| **Peak plantar pressure** | BM | 0.94 | -23.1 | 4.7e-8 |
| **Hindfoot / BM** | Neuromaturation | 0.39 | 6.9 | 0.22 |
| | **BM * Neuromaturation** | **0.99** | **-54.0** | **4.7e-11** |
| | BM * percent adult brain size | 0.98 | -38.9 | 8.7e-9 |
| **Peak plantar pressure** | BM | 0.92 | -21.6 | 1.6e-7 |
| **whole foot / BM** | Neuromaturation | 0.55 | 1.7 | 3.7e-3 |
| | **BM * Neuromaturation** | **0.99** | **-51.3** | **1.9e-10** |
| | BM * percent adult brain size | 0.99 | -47.9 | 6.2e-10 |

*SOM Table S.1. FTI/BM = force-time-integral normalized by body mass, BM = body mass (Newtons), S.2 Peak plantar pressure.*

Predictions of FTI/BM and peak plantar pressure data from the models are plotted in SOM Figure S.1. Based on these very strong correlations we infer that the interaction between neuromaturation (and by proxy, percent adult brain size) and body mass accurately predicts the force-time-integral (loading environment) of the foot, both peak plantar pressure in the whole foot and hindfoot, and body mass normalized force-time-integrals in the whole foot and hindfoot. These plots support our model where the loading environment of the foot can be accurately predicted by the interaction between neuromaturation (or by proxy, percent adult brain size) and body mass.

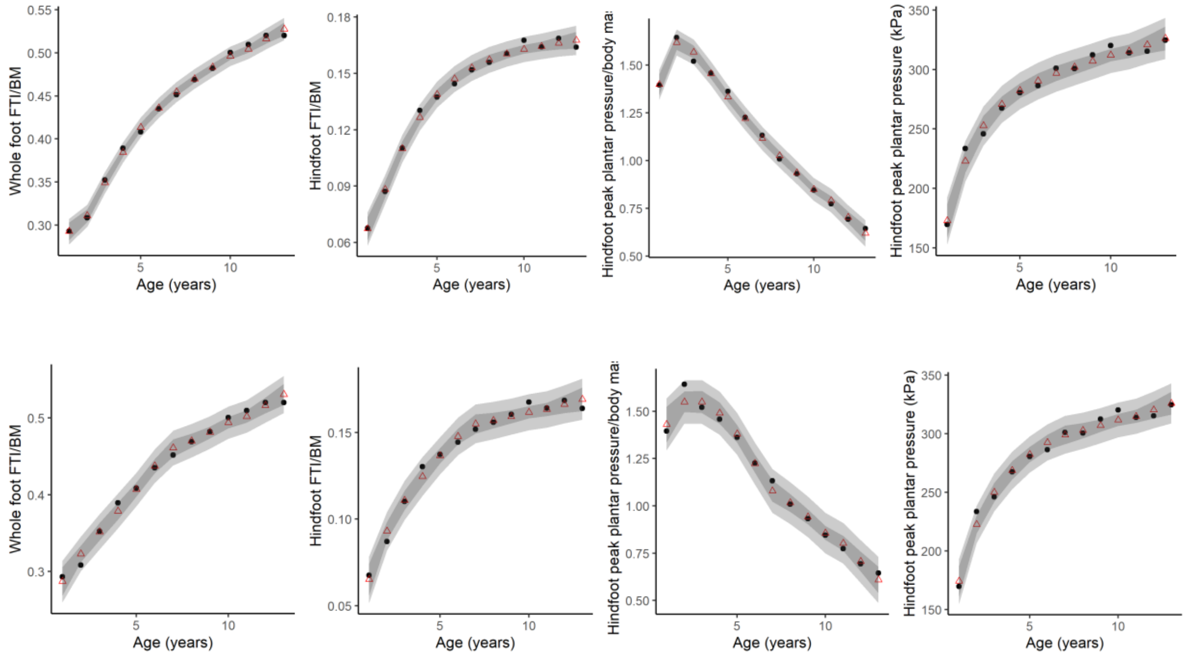


*SOM Figure S.1. (Top) foot loading variables with age predicted by interaction between body mass and neuromaturation. (Bottom) foot loading variables predicted by interaction between body mass and percentage of adult brain size.*

The force-time integral relative to body size (FTI/BM) with age can be almost perfectly predicted by the level of neural maturation (or percent brain size) and body size of an individual. This very strongly supports our model that neural maturation could be an important factor that contributes to changing loading patterns as gait develops. Furthermore, BV/TV in the human calcaneus [taken from Saers et al. (2020)] is predicted by the same interaction between body mass and neuromaturation (Adj. $R^2$=.85, p= 2.7e-5, see plot below). At least some of the error in BV/TV underlying the lower $R^2$ can be explained by the fact that BV/TV was measured in a small sample of archaeological specimens for which age and body size had to be estimated. Meanwhile, FTI and neuromaturation were calculated from equations based on mean trends in large samples from the literature, and therefore do not contain variation that would be associated with intrinsic individual differences and variation in the tempo at which individuals mature, which is there in BV/TV. Overall, this correlational evidence is highly encouraging and future experimental longitudinal study may be able to show the exact causal mechanism.

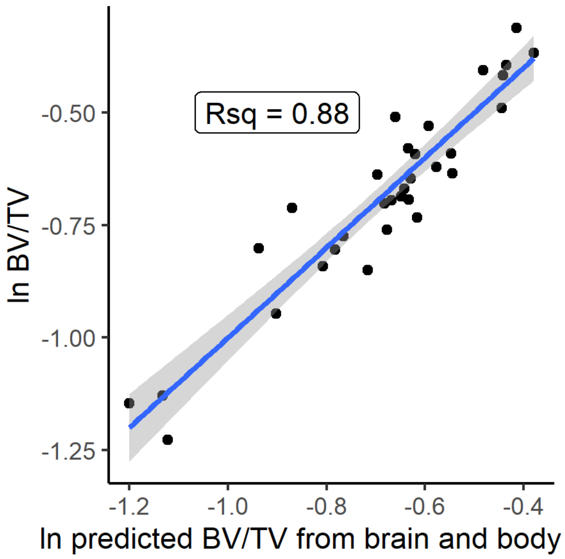


SOM Figure S.2. *Regression between measured ln BV/TV (from Saers et al. (2020)) and Bv/TV predicted by an interaction between neuromaturation and body size.*

### SOM References

Carter, D.R., Van Der Meulen, M.C.H., Beaupré, G.S., 1996. Mechanical factors in bone growth and development. Bone. 18, S5–S10.

Cofran, Z., Desilva, J.M., 2015. A neonatal perspective on Homo erectus brain growth. Journal of Human Evolution. 81, 41–47.

Fair, D.A., Schlaggar, B.L., 2008. Brain Development. Encyclopedia of Infant and Early Childhood Development.

Forssberg, H., 1985. Ontogeny of human locomotor control I. Infant stepping, supported locomotion and transition to independent locomotion. Experimental Brain Research. 57, 480–493.

Forssberg, H., 1999. Neural control of human motor development. Current Opinion in Neurobiology. 9, 676–682.

Müller, S., Carlsohn, A., Müller, J., Baur, H., Mayer, F., 2012. Static and dynamic foot characteristics in children aged 1-13 years: A cross-sectional study. Gait and Posture. 35, 389–394.

Okamoto, T., Okamoto, K., Andrew, P.D., 2003. Electromyographic developmental changes in one individual from newborn stepping to mature walking. Gait and Posture. 17, 18–27.

Saers, J.P.P., Ryan, T.M., Stock, J.T., 2020. Baby steps towards linking calcaneal trabecular bone ontogeny and the development of bipedal human gait. Journal of Anatomy. 236, 474–492.

Sutherland, D., 1997. The development of mature gait. Gait & Posture. 6, 163–170.

Vaughan, C.L., Langerak, N.G., OMalley, M.J., 2003. Neuromaturation of human locomotion revealed by non-dimensional scaling. Experimental Brain Research. 153, 123–127.
